# Characterization of neuronal ensembles in a model of dual reward conditioned place preference

**DOI:** 10.1101/2025.06.11.659163

**Authors:** Levi T. Flom, Kathryn L. Sandum, Skylar L. Hodgins, Samuel Johnson Noya, Jenna Crouse, Zhaojie Zhang, Ana-Clara Bobadilla

## Abstract

Substance use disorder (SUD) is associated with maladaptive alterations in behavior. Drug-seeking behavior has been associated with neuronal ensembles, defined here as a small group of neurons exhibiting coordinated activity patterns, in the prelimbic prefrontal cortex (PL) and nucleus accumbens core (NAcore). Most SUD preclinical research focuses on the effects of single-reward exposure on the general population of neurons within the reward pathway, rather than on poly-reward exposure. Here, we seek to characterize and compare the ensembles linked to cocaine and chocolate using a within-subject approach. We used Ai14xFos^2A-iCreER^ (c-Fos-TRAP2) transgenic mice to tag neuronal ensembles in a novel dual cocaine and chocolate conditioned place preference (CPP) model, where each chamber was associated with a different reward, either cocaine or chocolate. We found that, after successful dual conditioning and in the absence of the rewards, mice preferred cocaine over chocolate. Additionally, in mice exposed to both cocaine and chocolate, cortical and accumbal ensembles linked to each reward were comparable in size to reward ensembles in mice exposed only to one reward. However, reward-seeking ensembles were larger than ensembles tagged in homecage control mice across groups. We also found that ensemble size does not correlate with the level of reward seeking across single and dual CPP models. These results offer a new paradigm for studying drug-related neuronal ensembles in comparison to natural rewards in non-contingent behavioral models.

## Introduction

Substance use disorder (SUD) is characterized by uncontrolled use of a substance despite adverse consequences, often failing to meet social, financial, and personal responsibilities (1). In this disorder, drug rewards are prioritized over natural rewards, such as food. Both overlapping and specific neural circuitry and behavioral outcomes drive the seeking of natural and drug rewards (2). To understand the specific changes that occur while seeking drug rewards, we crossed Ai14 x Fos^2A-iCreERT2^ (c-Fos-TRAP2) transgenic mice (3) to tag reward-specific neuronal ensembles, defined here as co-active subpopulations of cells tied to the specific behavior of reward seeking (4). Commonly used markers to tag ensemble cells during behavioral events include immediate early genes, such as cellular oncogene FOS (c-Fos) (5) and activity-regulated cytoskeleton-associated protein (Arc) (6); other techniques focused on neuronal firing via calcium signaling also provide valuable insights into neuronal ensemble formation and specificity (7,8). Much research has focused on various learning experiences to characterize neuronal ensembles, including fear conditioning and SUD (9–13). Neuronal ensembles play a critical role in understanding the causes of maladaptive behaviors associated with SUD, some of which are absent in response to natural rewards (14).

While the study of the exposure to single drugs in animals has brought invaluable knowledge on the neurobiology of the reward circuitry, humans struggling with SUD often use multiple drugs and natural rewards concurrently (15). Preclinical rodent research on polysubstance SUD is typically conducted in rats, usually comparing two drugs of abuse as rewards (16–18). Here, we aim to expand research on both poly-reward exposure and ensemble specificity (19–21) in response to cocaine and chocolate rewards in mice. Another key aspect of modeling SUD is the consideration of both contingent and non-contingent models, each of which has its merits (22). Contingent models, such as self-administration (SA), facilitate the learning and reinforcement of actions leading to reward acquisition but require more time for associations to form. Non-contingent models, such as conditioned place preference (CPP), have faster experimental timelines but sacrifice some specificity due to reduced volitional control over the exposure to rewards. Research on poly-reward SA for cocaine and sucrose has shown that neuronal ensembles for different rewards are distinct (11,20). However, this distinction is difficult to test in non-contingent models due to the lack of operant manipulation to induce specific reward-seeking behavior on test day. Here, we present a CPP model in which each CPP chamber is associated with a distinct reward. Mice experience both drug (cocaine) and natural (chocolate) rewards in the two distinct contexts. We measured reward preference during testing based on the time spent in each chamber. This design enables a direct comparison of the gratifying properties of each reward.

The goal of this study is to use a non-contingent dual model to tag and measure the size of cocaine and chocolate reward ensembles in the nucleus accumbens core (NAcore) and the prelimbic region (PL) of the prefrontal cortex. We focused on these two regions because the NAcore plays a significant role in cocaine use in mice, functioning as an integration center for inputs from the mesolimbic dopamine pathways (23,24). Additionally, PL outputs are necessary for social reward-seeking and cocaine-seeking (25,26). We hypothesize that cocaine and chocolate seeking within this dual CPP is associated with NAcore and PL ensembles. We also compared ensemble sizes in traditional single-reward CPP and examined the correlations between ensemble sizes and reward-seeking behaviors.

## Methods

### Animals

All animal procedures were performed in accordance with the University of Wyoming’s IACUC regulations. 7-24 week-old male and female mice were obtained by crossing female Ai14 knock-in mice (B6;129S6-Gt(ROSA)26Sortm14(CAG-tdTomato)Hze/J, stock # 007914, the Jackson Laboratory, Bar Harbor, ME, USA) with c-Fos-TRAP2 male knock-in mice (STOCK Fostm2.1(icre/ERT2)Luo/J, stock #030323, the Jackson Laboratory). Mice were individually housed on a reverse 12-hour light cycle at least 3 days prior to the start of the experiment. Mice involved in chocolate or dual-conditioned place preference (CPP) experiments were given access to 3-7 hours of *ad libitum* food every day starting from the preconditioning day, except for mice that lost more than 20% of their initial weight or appeared unwell. Mice involved in single cocaine CPP were given *ad libitum* food access. Mice involved in chocolate and dual CPP experiments were given ten miniature chocolate chips (Nestle, Vevey, Switzerland) in their homecage the day before preconditioning started to prevent neophobia. Mice were handled and received at least 3 vehicle intraperitoneal (IP) injections over 3 days to acclimate them to handling.

### Conditioned Place Preference (CPP)

A 3-chamber setup (Med Associates, Fairfax, VT, USA) with a white chamber with mesh flooring, a black chamber with rod flooring, and a gray connecting chamber was used for CPP experiments. On preconditioning days, mice were placed in the grey center chamber and allowed to move freely through all three chambers for 30 minutes. Single-reward CPP conditioning followed a biased design, where the least preferred compartment for each subject was assigned as the reward-paired compartment (chocolate for chocolate CPP experiments and cocaine for cocaine CPP experiments). The assignment was counterbalanced for dual CPP experiments. On conditioning days, mice were confined to the less preferred chamber for 15 minutes with chocolate or cocaine administration and to the preferred chamber with vehicle (saline) on alternating days. This continued for four conditioning sessions with the reward and four with the vehicle, as shown in **Figure 1A**. Mice were given vehicle injections before being placed into the chamber for experiments that involved cocaine on non-cocaine days and were injected with 10mg/kg of cocaine (NIDA Drug Supply Program, Bethesda, MD) on cocaine-conditioning days. Ten miniature chocolate chips (Nestle, Vevey, Switzerland) were placed into a trough (½ of an empty 50 mL Falcon conical tube) (Corning, Corning, NY, USA) that was secured to the floor of the chocolate-paired chamber on chocolate conditioning days and an empty trough was secured to the non-chocolate-paired chamber on vehicle conditioning days. An additional “saline CPP” group was included to control for the experimental environment’s impact on c-Fos expression, where mice in both chambers were given saline injections across conditioning days. On test days, mice were placed into the gray center chamber and allowed to move freely through all three chambers for 30 minutes. Immediately after the 30 minutes, mice were given IP injections of 50 mg/kg 4-hydroxytamoxifen (4-OHT) (Sigma Aldrich, St. Louis, MO, USA) prepared per previous literature (3) Briefly, 10 mg of 4-OHT powder was dissolved in 250 mL of dimethyl sulfoxide (DMSO, Biorad, Hercules, CA USA) and frozen. Each of these aliquots was brought to room temperature and then added to 400 μL of 25% Tween 80 (Sigma-Aldrich) and 4.35 mL of saline, followed by vigorous vortexing until clear, 5 minutes before the conclusion of the test day. Additionally, a control group underwent handling and saline IP injections for three days before receiving an injection of 50mg/kg 4-OHT in the homecage to control for basal expression of c-Fos.

**Figure 1.**
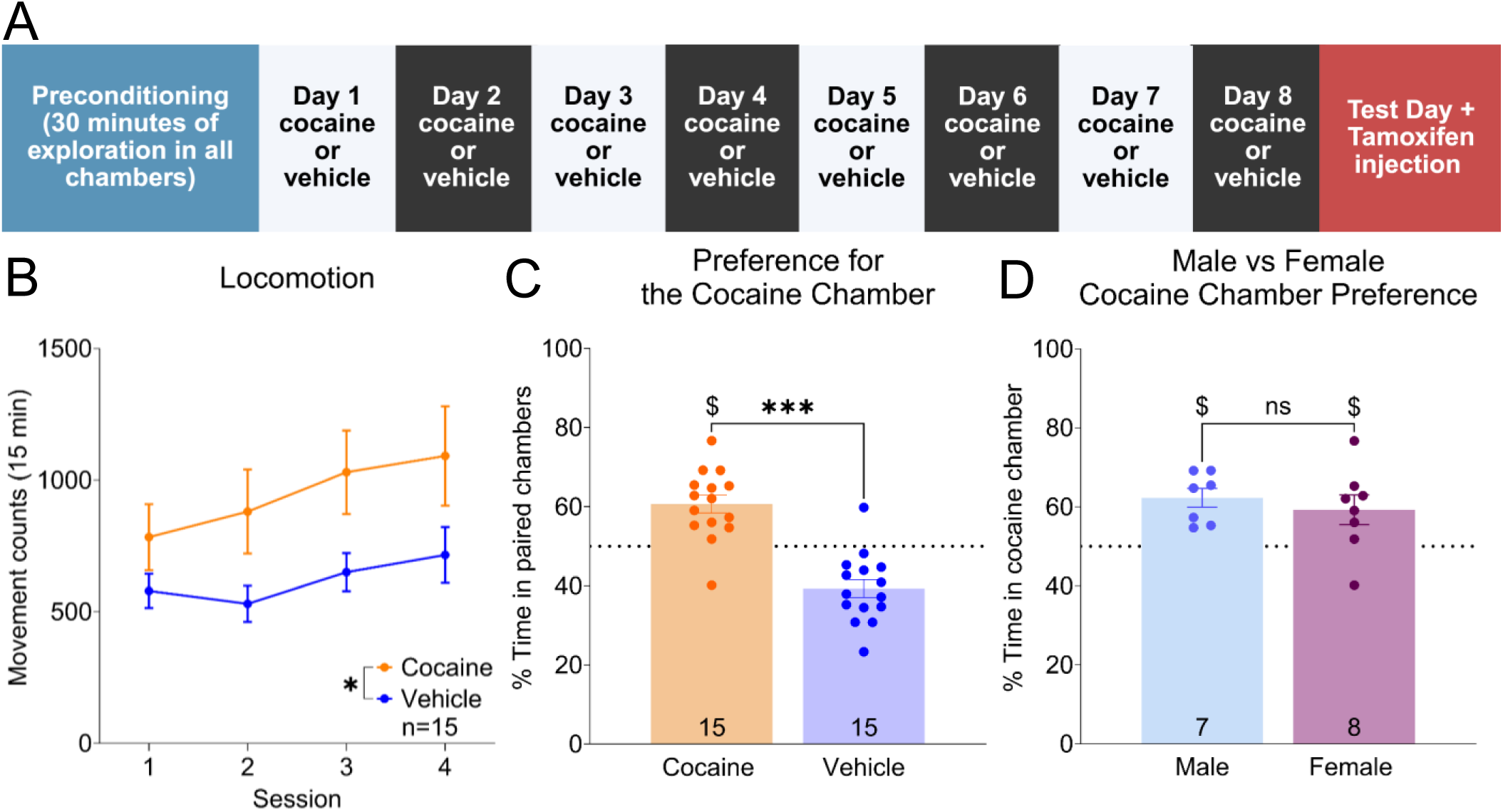
Behavior of Cocaine CPP. (**A**) Timeline of cocaine CPP. Different colors denote different contexts. (**B**) Movement counts during conditioning sessions for the cocaine-paired chamber and the vehicle-saline chamber. * p < 0.05 Comparing movement counts during conditioning. (**C**) Percent time in each chamber on test day. The dotted line signals 50% time, or no preference. *** p < 0.001 Comparing time in the cocaine-paired chamber to the vehicle-saline chamber, paired t-test. $ p < 0.05 Cocaine compared to 50%, one-sample t-test. (**D**) No sex differences in cocaine CPP. $ p < 0.05, Males and females compared to 50%, one-sample t-test. Numbers at the bottom of the bars refer to the number of animals.

### Immunohistochemistry

Mice were anesthetized with a 2:1 combination of ketamine (MWI, Boise, ID, USA) and xylazine (Akorn, Lake Forest, IL, USA), then perfused with phosphate-buffered saline (PBS, Quality Biological, Gaithersburg, MD, USA) and 3.7% formaldehyde (Fisher Scientific, Fairlawn, NJ, USA). Brains were post-fixed in 3.7% formaldehyde for at least 24 hours and then transferred to a solution of 20% sucrose + 0.01% sodium azide (Sigma-Aldrich) in 1X PBS. 50-μm brain sections were cut using a cryostat (Leica, Deer Park, IL, USA) from approximately Bregma 2.0 to Bregma 0.62. Free-floating sections were rinsed 3 times in 1X PBS and incubated for 2 hours in a blocking solution of 5% normal goat serum (NGS, Invitrogen, Waltham, MA USA), 2.5% bovine serum albumin (BSA, Fisher Scientific), and 0.25% Triton x 100 (Thermo Scientific, Waltham, MA USA) in PBS, then incubated in this blocking solution plus the primary antibody for 16-24 hours. Anti-Neuronal Nuclei (NeuN) primary antibody (mouse, 1:1000, EMD Millipore Core, Burlington, MA, USA) was used for ensemble quantification. The tissue was then rinsed 1 time in 0.25% PBS-Triton, 3 times in 1X PBS, and incubated for 2 hours in secondary antibodies. We used a 1:1000 goat anti-mouse IgG (H+L) cross-adsorbed secondary antibody, Alexa Fluor™ 647 (Invitrogen, Thermo Fisher, Waltham, MA, USA) for ensemble quantification. The tissue was mounted with Prolong Gold (Life Technologies Corporation, Carlsbad, CA, USA).

### Image Acquisition and Quantification

Images of the NAcore with the anterior commissure and PL were taken using a 63x, 1.4 NA, oil immersion objective on a Zeiss 980 confocal microscope (Zeiss, Oberkochen, Germany) at the Integrated Microscopy Core, University of Wyoming, or a Zeiss 880 confocal microscope (Zeiss) at Colorado State University. The laser lines used were green (555 nm) for TdTomato and far red (639 nm) for NeuN. For each mouse, 4 to 6 areas of both the NAcore and PL were randomly selected and imaged. Each image was a 3 x 3 tile (512 x 512 pixels per image, 380 μm^2^), and a Z-stack with a 1 μm step size was acquired, comprising 16-25 images per stack. Images were then stitched in Zeiss Blue software (Zeiss) and converted to Imaris software (Oxford Instruments, United Kingdom) for image analysis. Before analysis, images were cropped to a standard 15 μm to ensure uniform image thickness. The NeuN cell bodies (approximately 7 μm in diameter) were semi-automatically counted using the “spots” function and the mean intensity of the channel. A blinded experimenter manually checked and adjusted all automated counts. The tdTomato+ cell bodies (about 10 μm in diameter) were manually counted. TdTomato+ percent expression was established by dividing the tdTomato+, NeuN+ overlap cells by the total NeuN+ in the images. One animal was removed from the single-cocaine experiment due to the lack of expression of TdTomato+ cells following conditioning.

### Statistical analysis

Statistics were performed using Prism 10 (GraphPad Software, CA, USA). Behavioral results were analyzed using two-way analysis of variance (ANOVA) or paired t-tests, as appropriate. One-sample t-tests compared to 50% validated successful conditioning, not due to chance. Ensemble size in the regions of interest was compared with a paired t-test to account for individual differences between mice. Between-reward analysis of ensemble quantification was performed using a two-way ANOVA. Post-hoc analysis was completed using Tukey’s multiple comparison tests. Ensemble correlation with preference was performed using regression analysis. Significance was set to p < 0.05, and all data are presented as mean ± SEM. Outliers were tested for using ROUT and a Q of 0.5%. One mouse was removed from the single-reward cocaine ensemble group following outlier testing.

## Results

### Single Cocaine CPP Behavior

We determined the ensemble size for cocaine-seeking behavior in a standard single-reward CPP (**Figure 1A**). As expected, c-Fos-TRAP2 mice showed a significant increase in locomotion in the cocaine-paired chamber (**Figure 1B**, two-way ANOVA, F (1,29) = 4.561, p = 0.0413, 95% CI = 13.88-641.8). On test day, mice showed a significant preference for the cocaine-paired chamber compared to the chamber conditioned with saline vehicle (**Figure 1C**, paired t-test, t14 = 4.729, p = 0.0003, 95% CI = 11.66-31.00, cocaine chamber 60.7% ± 2.26, vehicle chamber 39.34% ± 2.26). The preference for the cocaine-paired chamber was also significantly higher than chance (one-sample t-test compared to 50%, t14 =4.729, p = 0.0003, 95% CI = 5.828-15.50). No sex differences were observed (**Figure 1D**, unpaired t-test, n_male_ = 7, n_female_ = 8, t13 = 0.6582, p = 0.5219, 95% CI = −13.01-6.931, male 62.28% ± 2.41, female 59.25% ± 3.76). Both sexes showed an increase in the cocaine-paired chamber compared to chance (**Figure 1D**, One-sample t-tests compared to 50%, n_male_ = 7, t6 = 5.102, p = 0.0022, 95% CI = 6.392-18.18, n_female_ = 8, t7 = 2.462, p = 0.0433, 95% CI = 0.3664-18.13).

### Single Chocolate CPP Behavior

A total of 19 c-Fos-TRAP2 mice trained in the single chocolate CPP (**Figure 2A**) showed no difference in locomotion during conditioning (**Figure 2B**, two-way ANOVA, F (1,20) = 0.5528, p = .4658, 95% CI = −233.3-106.0). Following conditioning, mice significantly preferred the chocolate-paired chamber on test day compared to the chamber where chocolate was never presented during conditioning (**Figure 2C**, paired t-test, t18 = 2.885, p = 0.0099, 95% CI 4.609-29.32, chocolate chamber 58.48% ± 2.941, empty chamber 41.52% ± 2.941). Mice preferred the chocolate-paired chamber significantly more than chance (**Figure 2C**, one-sample t-test compared to 50%, t18 = 2.885, p = 0.0099, 95% CI = 2.305-14.66). Males showed a significantly higher preference for the chocolate-paired chamber than females (**Figure 2D**, unpaired t-test, n_male_ = 9, n_female_ = 10, t17 = 2.589, p = 0.0191, 95% CI = −24.12--2.461, male 65.48% ± 3.684 female 52.19% ±3.565) and only males showed a significantly higher preference for the chocolate-paired chamber compared to chance (**Figure 2D**, One sample t-tests compared to 50%, n_male_ = 9, t8 = 4.201, p = 0.0030, 95% CI = 6.982-23.97, n_female_ = 10, t9 = 0.6137, p = 0.0433, 95% CI = −5.877-10.25).

**Figure 2.**
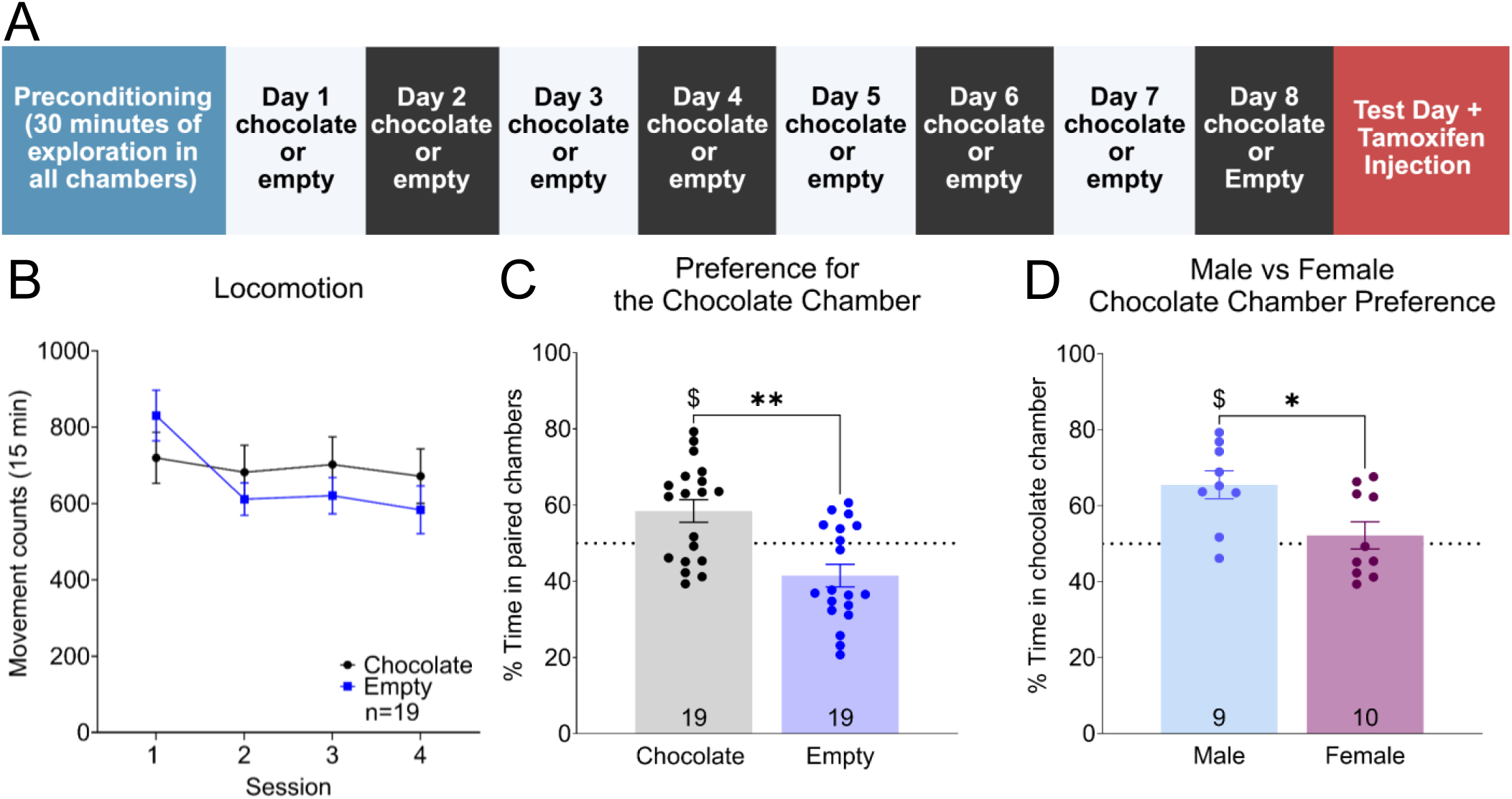
Behavior of Chocolate CPP. (**A**) Timeline of chocolate CPP. Different colors denote different contexts. (**B**) Movement counts during conditioning sessions for the chocolate-paired chamber and the vehicle-empty chamber. (**C**) Percent time in each chamber on test day. The dotted line signals 50% time, or no preference. ** p < 0.01 Comparing time in the chocolate-paired chamber to the empty-paired chamber, paired t-test. $ p < 0.05 Chocolate chamber compared to 50% one-sample t-test. (**D**) Sex differences in chocolate preference showed that male mice preferred chocolate more than females. * p < 0.05 Comparing chocolate chamber time between sexes, unpaired t-test. $ p < 0.05, Males compared to 50% one-sample t-test. Numbers at the bottom of the bars refer to the number of animals.

### Dual cocaine and chocolate CPP behavior

In our new paradigm of dual cocaine and chocolate CPP, we alternated cocaine and chocolate days in distinctive chambers during conditioning (**Figure 3A**). The c-Fos-TRAP2 mice showed significantly elevated locomotion during conditioning in the cocaine-paired chamber compared to the chocolate-paired chamber (**Figure 3B**, two-way ANOVA, F (1,20) = 30.65, p < 0.0001, 95% CI = 674.4-1490). On test day, mice significantly preferred the cocaine-paired chamber to the chocolate-paired chamber (**Figure 3C**, paired t-test, t10 = 2.953, p = 0.0145, 95% CI = −22.91--3.204, cocaine chamber 56.53% ± 2.21, chocolate chamber 43.47% ± 2.21) and the preference was significantly higher than chance (**Figure 3C**, one sample t-test compared to 50%, t10 = 2.953, p = 0.0145, 95% CI = 1.602-11.46). No sex differences were observed between groups (**Figure 3D**, unpaired t-test, n_male_ = 6, n_female_ = 5, t9 = 0.7733, p = 0.4592, male cocaine 54.94% ± 3.89, female cocaine 58.44% ±1.56). However, only females showed a preference for the cocaine-paired chamber higher than chance, while males did not (**Figure 3D**, one-sample t-tests compared to 50%, n_male_ = 6, t5 = 1.270, p = 0.2601, 95% CI = −5.058-14.93, n_female_ = 5, t4 = 5.291, p = 0.0061, 95% CI = 4.011-12.87).

**Figure 3.**
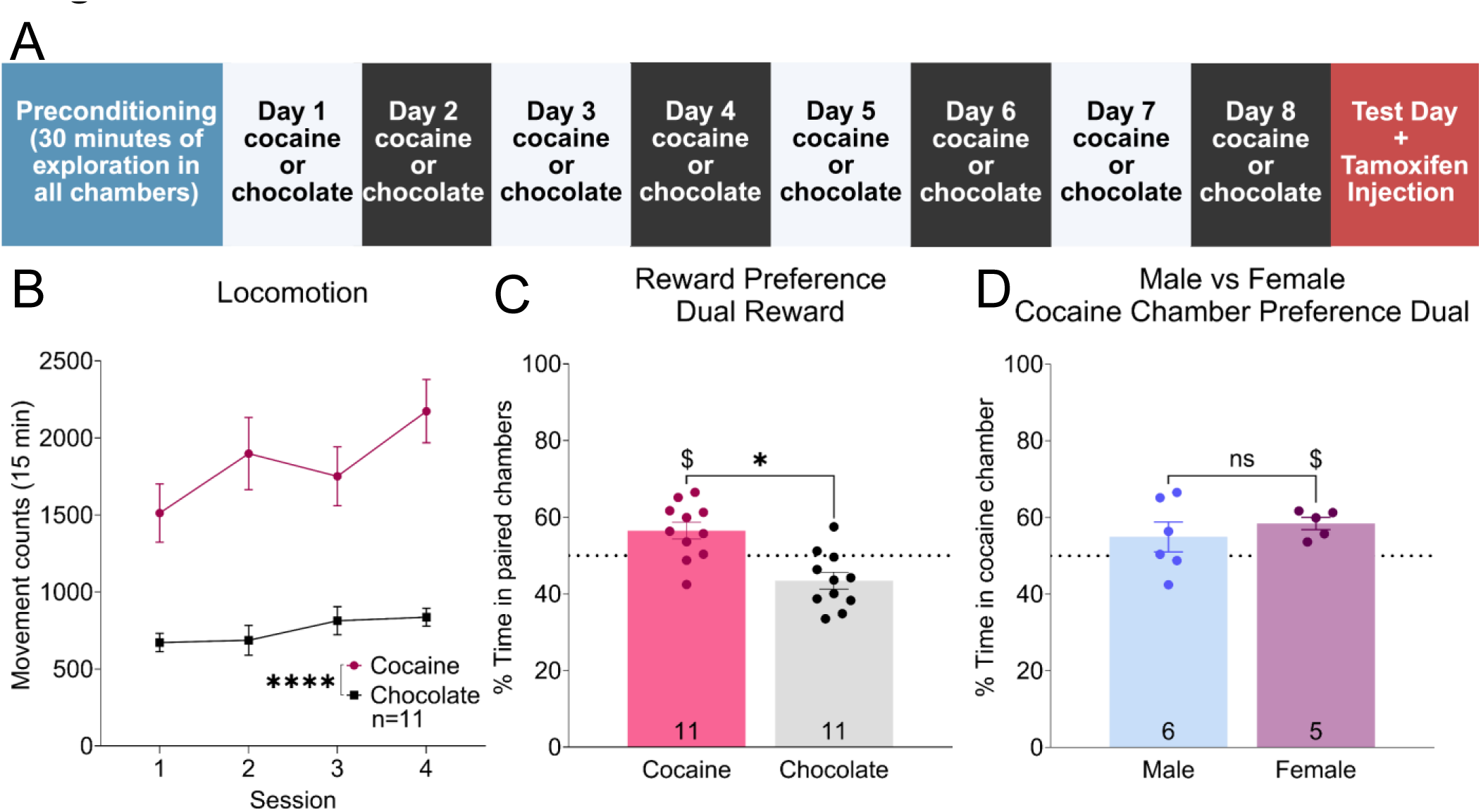
Behavior of dual CPP. (**A**) Timeline of dual CPP. Different colors denote different contexts. (**B**) Movement counts during conditioning sessions for the cocaine-paired chamber and the chocolate-paired chamber. **** p < 0.0001 Comparing movement counts during conditioning. (**C**) Percent time in each chamber on test day. The dotted line signals 50% time, or no preference. * p < 0.05 Comparing the cocaine-paired chamber and the chocolate-paired chamber, paired t-test. $ p < 0.05 Chamber preference compared to 50% one-sample t-test. (**D**) Male and female preferences for cocaine were not different from each other. $ p < 0.05 Female cocaine preference compared to 50% one-sample t-test. Numbers at the bottom of the bars refer to the number of animals.

### Ensemble characterization

To compare the ensemble sizes between regions and across different types of rewards, we quantified TdTomato and NeuN signals; see **Figure 4A** for a representative image in the NAcore and **Figure 4B** for a representative image in the PL. All neuronal ensembles were larger in the PL than the NAcore: cocaine-seeking ensemble in the single cocaine CPP (**Figure 4C**, paired t-test, t18 = 15.68, p < 0.001, 95% CI = 1.594:2.087, NAcore 0.65% ± .07, PL 2.5%), chocolate-seeking ensemble in the single chocolate CPP (paired t-test, t18 = 15.68, p < 0.001, 95% CI = 1.594-2.087, NAcore 0.65% ± 0.07, PL 2.5% ± 0.12), and cocaine-seeking ensemble in the dual CPP (paired t-test, t10 = 6.216, p < 0.0001, 95% CI = 1.101-2.332, NAcore 0.67% ± 0.15, PL 2.39% ± 0.27). To validate that the tagged ensembles above were associated with reward seeking, a control group of 14 c-Fos-TRAP2 mice underwent saline CPP in both chambers without exposure to any other reward and received a 4-OHT injection immediately following the test session (**Supplemental Figure S1A**). Assessing the locomotion of the animals during conditioning revealed a difference between the black and white chambers, with increased movement observed in the black chamber compared to the white (**Supplemental Figure S1B**, two-way ANOVA F (1,30 = 38.37, p < 0.0001, 95% CI = 230.9-458.0). A probable explanation for this is due to differences in the flooring, with mice preferring solid bar flooring over mesh (27). As expected, there was no preference between chambers on test day (**Supplemental Figure S1C**, paired t-test, t13 = 1.202, p = 0.2509, 95% CI = - 6.170-21.64, white chamber 53.87% ± 3.218, black chamber 46.13% ± 3.218). In evaluating sex differences on our test day, we show no difference in preference for the black chamber (**Supplemental Figure 1D**, unpaired t-test, n_male_ = 9, n_female_ = 5, t12 = 1.366, p = 0.1970, Effect size = −8.882, 95% CI −23.05-5.286, male 49.31% ± 13.58, female 40.42% ± 6.235). However, only females showed a decrease in time for the black chamber higher than chance, while males did not (**Supplemental Figure S1D**, one-sample t-test compared to 50% n_male_ = 9, t8 = 0.1534, p = 0.8819, 95% CI = −11.13-9.745, n_female_ = 5, t4 = 3.435, p = 0.0264, 95% CI = −17.32-1.836).

**Figure 4.**
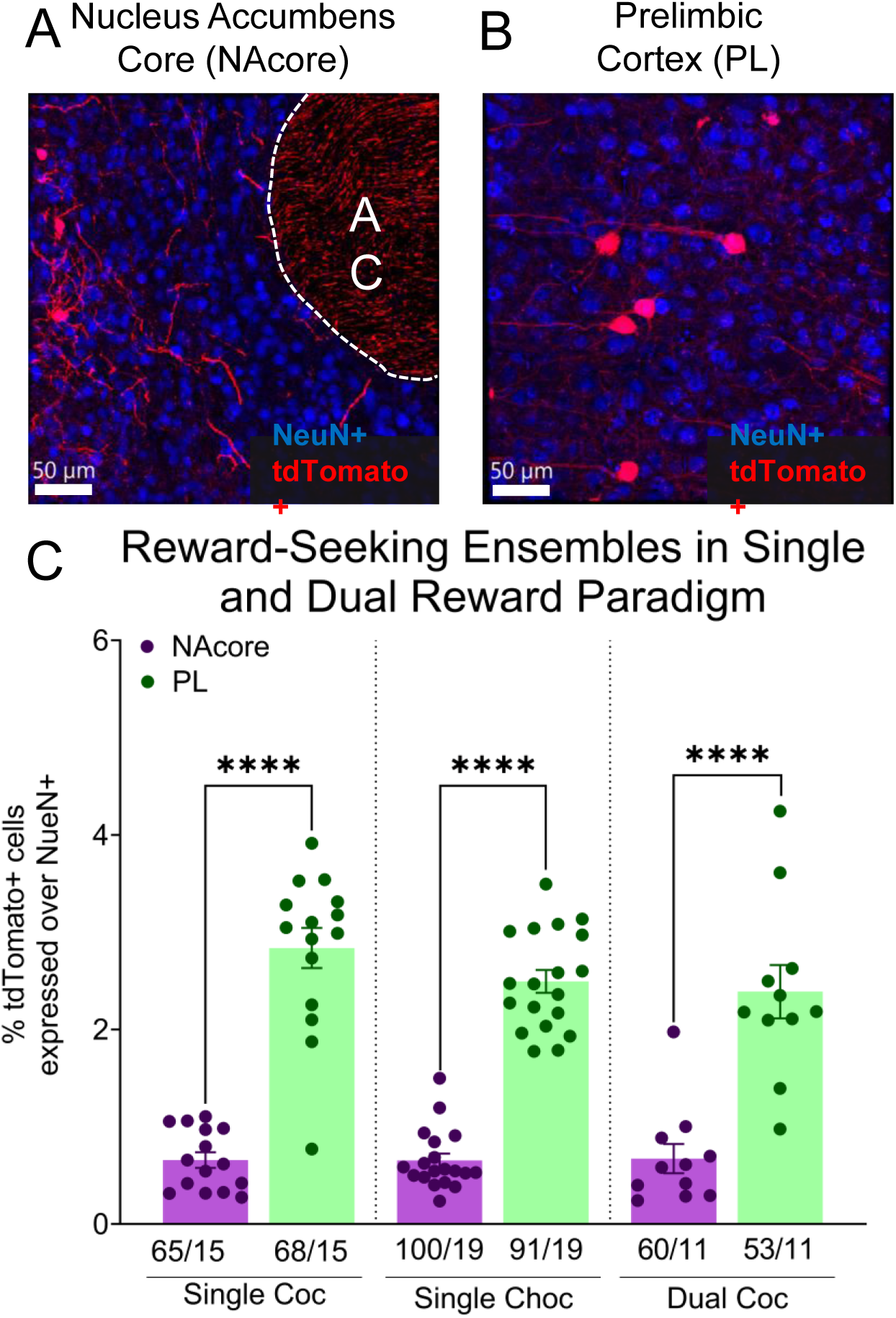
Ensemble sizes following single- and dual-reward CPP. (**A**) Representative image of the nucleus accumbens core (NAcore), Red TdTomato+, Blue NeuN+. (**B**) Representative image of the prelimbic cortex (PL). (**C**) Size of the reward-seeking ensemble in the NAcore and Prelimbic cortex across reward groups. Anterior commissure (AC), cocaine (Coc), chocolate (Choc), **** p < 0.0001. Numbers refer to the number of image acquisitions per animal.

When comparing the size of ensembles in the NAcore across groups, we observed a significant difference in ensemble size (**Figure 5A**, two-way ANOVA, F (4,279) = 2.999, p = 0.0190). A post-hoc analysis revealed that the saline ensemble, cocaine-seeking ensemble in the single cocaine CPP, chocolate-seeking ensemble in the single chocolate CPP, and chocolate-seeking ensemble in the dual CPP were significantly larger than the homecage ensemble (homecage vs. saline, p = 0.0260, 95% CI = 0.05254-1.293; homecage vs. cocaine, p = 0.0142, 95% CI = 0.09750-1.338; homecage vs. chocolate, p = 0.02090, 95% CI = 0.06807-1.293; and homecage vs. cocaine in dual reward conditioning, p = 0.0260, 95% CI 0.1532-1.392). No differences were seen in ensemble size between the three reward conditions (cocaine-seeking ensemble in the single cocaine CPP, chocolate-seeking ensemble in the single chocolate CPP, and cocaine-seeking ensemble in the dual CPP).

**Figure 5.**
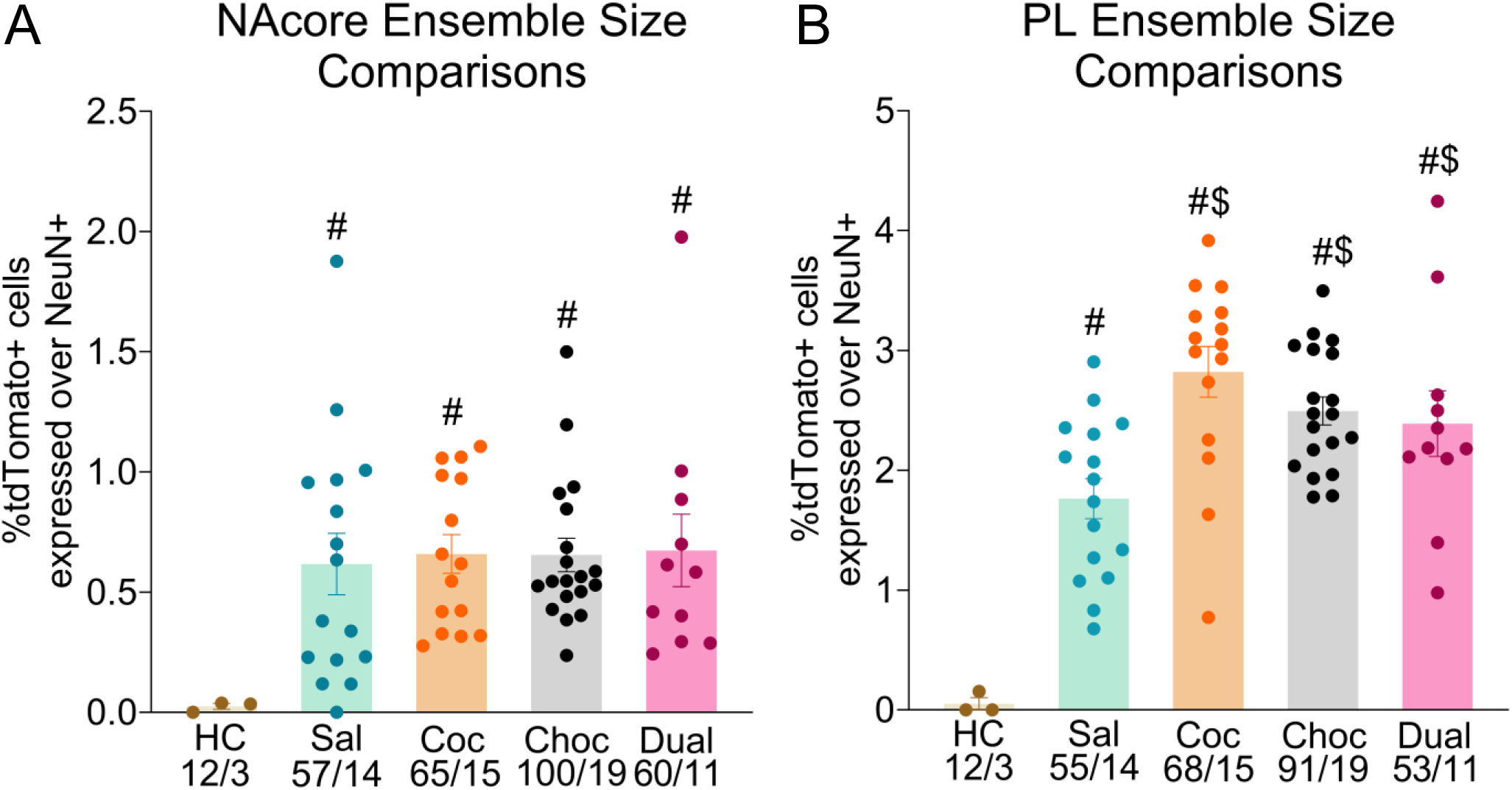
Comparison of ensemble sizes between groups. (**A**) Comparing the size of the ensemble in the different treatment groups in the NAcore. (**B**) Comparing the size of the ensemble in the PL. Nucleus Accumbens Core (NAcore), Prelimbic cortex (PL), homecage (HC), saline (Sal), cocaine (Coc), chocolate (Choc), dual cocaine (Dual), $ p < 0.05 compared to saline group in PL, # p < 0.0001 all groups compared to homecage group. Numbers refer to the number of image acquisitions per animal.

Comparisons of ensemble sizes in the PL revealed significant group differences (**Figure 5B**, two-way ANOVA, F (4,264) = 22.45, p < 0.0001). Post-hoc analysis showed that the homecage ensemble was significantly smaller than all other ensembles (saline, p < 0.0001, 95% CI = 1.074-3.010, cocaine in single CPP, p < 0.0001, 95% CI = 2.131-4.041, chocolate, p < 0.0001, 95% CI = 1.831:3.727, and cocaine in dual CPP, p < 0.0001, 95% CI =1.626-3.569). In contrast to the NAcore, the saline ensemble was also significantly smaller than reward-linked ensembles (cocaine in single CPP, p < 0.0001, 95% CI = 0.5294-1.560; chocolate, p < 0.0001, 95% CI = 0.2544-1.221; cocaine in dual CPP, p = 0.0494, 95% CI = 0.0008757-1.111). There was no difference in ensemble size between ensembles linked to the 3 reward conditions (cocaine-seeking ensemble in the single cocaine CPP, chocolate-seeking ensemble in the single chocolate CPP, and cocaine-seeking ensemble in the dual CPP).

To assess whether the size of the NAcore and PL ensembles relates to the seeking behavior found in our mice, we performed simple linear regressions of reward-seeking behavior on ensemble formation. We found no significance for either NAcore or PL in cocaine-seeking (NAcore: **Supplemental Figure S2A**, Simple linear regression, R2= .09030, p = 0.2765, PL: **Supplemental Figure S2B**, Simple linear regression, R2= 0.1233, p = 0.1994), chocolate seeking (NAcore: **Supplemental Figure S2C**, Simple linear regression, R2 = 0.02373, p = 0.5289, PL: **Supplemental Figure S2D**, Simple linear regression, R2 = 0.005476, p = .7634) and cocaine seeking in dual reward conditioning (NAcore: **Supplemental Figure S2E**, Simple linear regression, R2 = 0.2711, p = 0.1006, PL: **Supplemental Figure S2F**, Simple linear regression, R2 = 0.01307, p = 0.7378). These results show no correlation between ensemble size and reward-seeking behavior.

## Discussion

We developed a dual reward paradigm for neuronal ensembles tagging within a non-contingent dual model. We hypothesized that ensemble formation would occur in the NAcore and PL, with ensemble size correlating with reward-seeking behavior. Mice exhibited a preference for the cocaine- and chocolate-paired chambers following single-reward conditioning. This finding aligns with previous work demonstrating that non-contingent models effectively assess preference from rewarding stimuli (28,29). When presented with conditioning for both cocaine and chocolate, mice developed a preference for cocaine, demonstrating that cocaine reward supersedes natural reward seeking. We examined the size of neuronal ensembles associated with cocaine- and chocolate-seeking behaviors in the NAcore and PL following single and dual-reward CPP training. We observed a consistent ensemble size within the same region across rewards, and this size was not correlated with the seeking behavior.

While we observed no sex differences in the groups undergoing single cocaine CPP, males exhibited a significantly stronger preference for chocolate in the single chocolate CPP compared to females (**Figure 2D**). This finding contradicts previous reports of higher consumption of highly palatable food in female rats compared to male rats under low-cost conditions, with no differences in consumption observed at higher costs (30). In this context, “low cost” refers to conditions where access to food requires minimal effort, while “high cost” refers to conditions that require increased effort. This study by Freeman et al. utilized a within-session behavioral-economic paradigm, a contingent model in which active effort is required to receive food. In contrast, our approach employs a non-contingent model where food is provided independently of effort. These methodological differences may account for discrepancies in consumption across studies.

In the dual cocaine and chocolate CPP, we found that mice developed a preference for the cocaine-paired chamber, highlighting the elevated motivational value of cocaine cues relative to natural rewards when only the cues associated with rewards are present, but actual rewards are absent (31). This contradicts paradigms in which rats overwhelmingly chose non-drug sweetened water over cocaine when the actual rewards are delivered after the choice (32). However, this preference was later attributed to the different dopamine release pharmacokinetics between rewards (33,34). While we observed no significant differences between male and female mice in the dual cocaine and chocolate CPP, only females exhibited a preference for the cocaine-paired chamber above chance levels, whereas males did not (**Figure 3D**). This result aligns with previous findings that report a preference for the cocaine-associated cue over the sucrose-associated cue during cue-induced reinstatement in a mouse-contingent model of dual cocaine and sucrose self-administration (11). In both contingent and non-contingent dual reward paradigms, we conclude that the drug-associated cue or context holds greater motivational value than the non-drug reward when actual rewards are absent.

Regarding reward-associated neuronal ensembles, the most striking observation is that ensemble sizes in the PL are larger than those in the NAcore across single and dual cocaine and chocolate CPP paradigms (**Figure 4C**). This finding is consistent with previous literature (20,35) showing that the PL has higher levels of c-Fos expression following reward-seeking behaviors. This may be explained by a stronger expression of c-Fos in the PL compared to the NAcore (36), which would bias the ensemble tagging mechanism through c-Fos-TRAP2. We predict that the larger reward-linked ensemble in the PL could also correspond to a stronger reward-associated representation, given this region’s importance in decision-making, particularly during abstinence from cocaine and other drugs of abuse (37,38).

We also found that saline-, cocaine-, and chocolate-seeking ensembles are similar in size in the NAcore, regardless of whether single- or dual-reward paradigms were used. Comparable ensemble sizes in the NAcore associated with different rewards have also been observed in a contingent model of cocaine and sucrose self-administration (11). Surprisingly, despite mice not developing a preference for either saline-paired chamber (**Supplemental Figure 1C**), we observed a saline-associated ensemble in the NAcore that was equivalent in size to the reward-associated ensembles (**Figure 5A**). We hypothesize that the saline-associated ensemble is linked to the experimental context, specifically the different chambers within the CPP apparatus. This hypothesis is supported by the lack of neuronal ensembles formed in the homecage control group (**Figure 5**). This observation highlights the importance of context in both contingent and non-contingent models of reward research, as evidenced by the increase in c-Fos expression in response to a distinct context (CPP chambers) compared to a familiar one (Homecage). By observing an increase in c-Fos in both context-only and context-with-rewards, we believe that ensembles for rewards are also ensembles for the context the reward is paired with. Interestingly, while a saline-context ensemble was observed in the PL, it was significantly larger than the homecage group, yet smaller than the reward-seeking ensembles, regardless of whether single- or dual-reward exposure was used (**Figure 5B**). This suggests that ensemble size may distinguish between rewarding and non-rewarding contexts in the PL following repeated exposure. This finding contradicts previous work that showed that acute exposure to cocaine increased c-Fos expression in the NAcore but not the PL (39). This may be because a single exposure to cocaine is insufficient to induce high levels of c-Fos. In contrast, repeated exposure to contexts or stimuli in the PL produces stronger activation (3).

We employed c-Fos-TRAP2 transgenic mice (3) in our tagging method, which leverages c-Fos expression as a proxy for neuronal activation. It is essential to acknowledge the temporal limitations of this technique, as other methods have leveraged calcium-based activity-dependent tools with shorter tagging windows (40,41). However, our findings on ensemble size align with previous research across different drugs of abuse, underscoring the importance of c-Fos-expressing neuronal ensembles (4,19,42,43).

In this study, we did not find any correlation between ensemble size and cocaine-seeking behavior in CPP (**Supplemental Figure 2**). The results of our non-contingent protocol contrast with contingent protocols, where significant correlations have been found between the size of c-Fos-TRAP-tagged ensembles in the NAcore and the level of cocaine-seeking behavior after single- and dual-reward SA (11). Studies on ensembles using c-Fos-TRAP2 in fear recall suggest that learning duration (3) or the level of activation of individual cells during memory recall (44–47) may be stronger indicators of the role of ensembles on operant behavior. A similar interpretation was proposed in recent work showing that repeated non-contingent cocaine exposure reduced ensemble size while increasing ensemble signaling strength (47). This suggests that ensemble synaptic connective strength may be more important than size. One limitation of CPP is the reduced effort required from animals to learn reward-associated behaviors compared to operant conditioning (48). Although self-administration protocols often yield overlapping findings with CPP (49), motivational differences persist, highlighting the importance of both models in studying distinct phases of SUD (22). It is important to emphasize that only the groups that underwent the CPP protocols established ensembles linked to the context and rewards in the NAcore (**Figure 5A**). Additionally, ensembles associated with drug and non-drug rewards were significantly larger in the PL than those associated with context alone (saline CPP group) or homecage control groups (**Figure 5B**). Thus, in this non-operant model, PL ensemble size varies across different behavioral conditions, suggesting that size may be a distinguishing characteristic.

### Conclusions

We developed a novel non-contingent dual cocaine and chocolate CPP model, enabling us to characterize and compare drug- and non-drug-associated ensemble formation following poly-reward exposure to single-reward exposure. Given the growing interest in ensemble-centric studies within SUD research, characterizing neuronal ensembles in preclinical models of poly-reward exposure with strong face validity is crucial for advancing our understanding of SUD complexities and identifying potential therapeutic targets.

## Data Availability Statement

The raw and analyzed data that support the findings of this study are available from the corresponding author upon reasonable request.

## Acknowledgments

The authors thank the NIDA Drug Supply Program, administered by the Division of Therapeutics and Medical Consequences, for providing cocaine. They also thank the University of Wyoming’s Integrated Microscopy Core (IMC) for their technical assistance and all members of the Bobadilla Lab for their help and support.

## Supplemental Figures

**Supplemental Figure 1.**
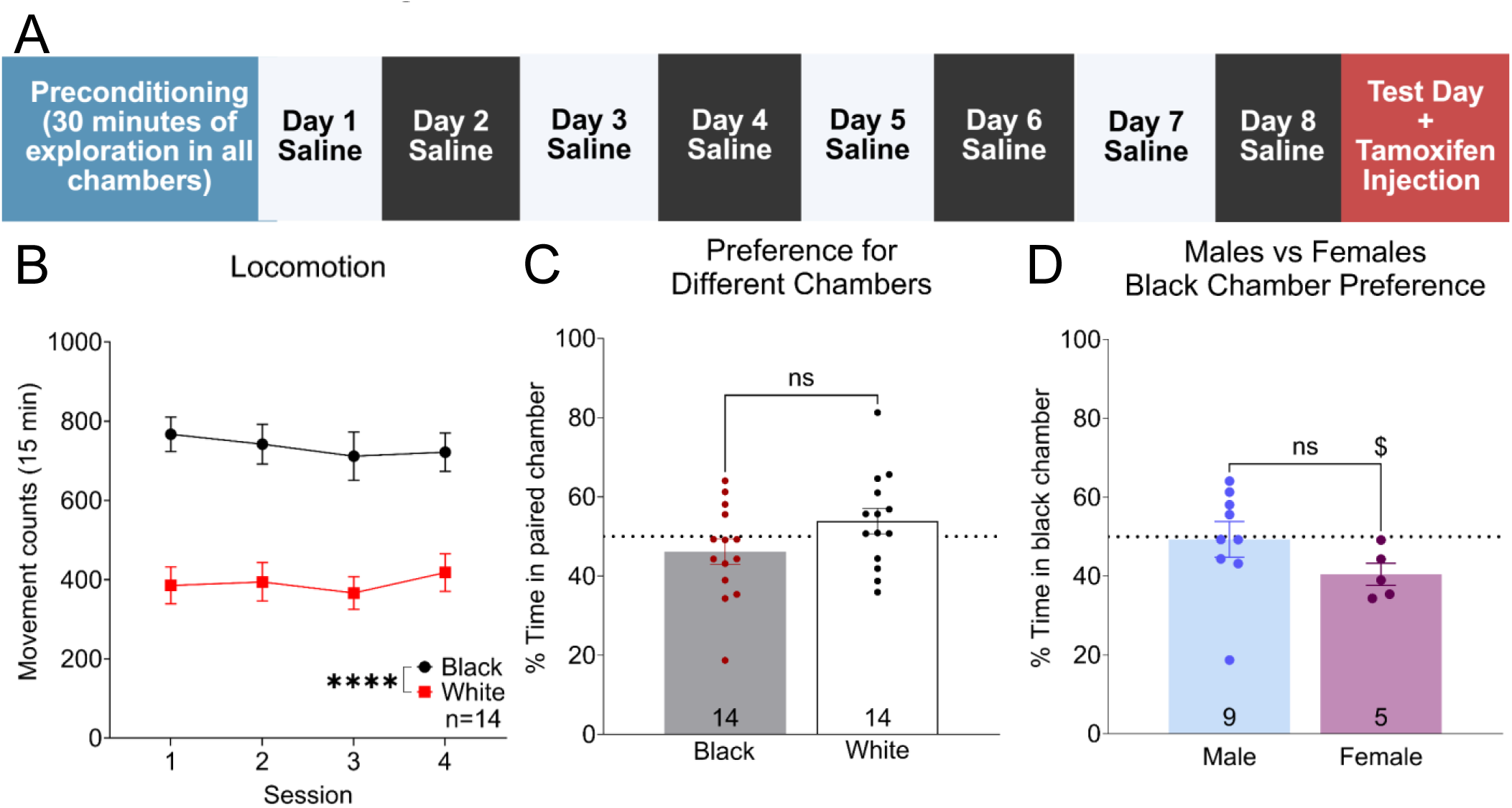
Behavior of Saline CPP. (**A**) Timeline of saline CPP. Different colors denote different contexts. (**B**) Movement counts during conditioning sessions for both black and white sessions. **** p < 0.0001 Comparing black and white chamber movement conditioning. (**C**) Percent preference shown for the different contexts on test day. (**D**) Sex differences in context preference were not observed, but females showed a reduction that differed from random chance. $ p < 0.05 black chamber preference for females compared to 50%. Numbers at the bottom of the bars refer to the number of animals.

**Supplemental Figure 2.**
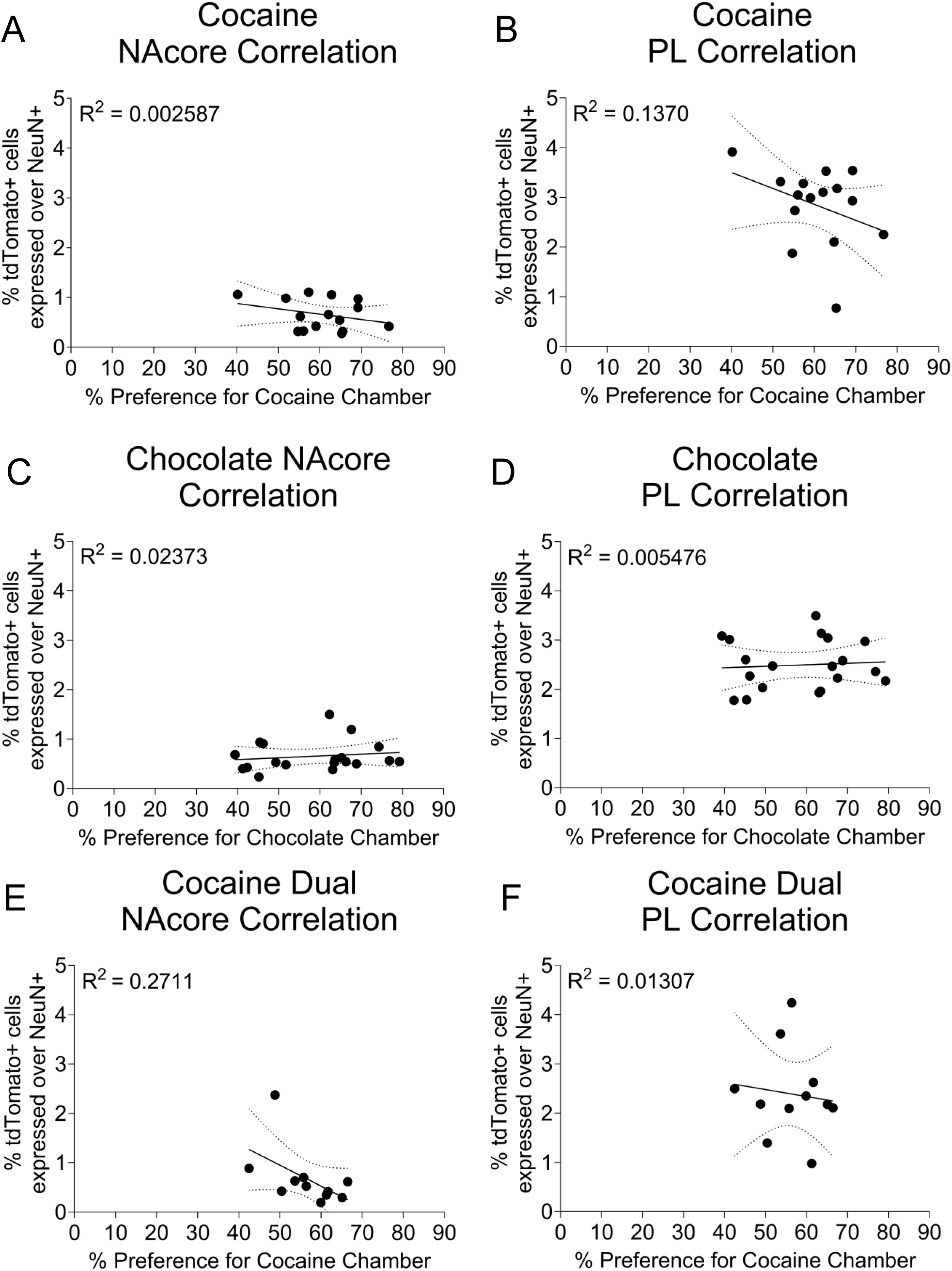
Ensemble size does not correlate with seeking levels. (**A**) Correlation of cocaine seeking with the NAcore ensemble size. (**B**) Correlation of cocaine seeking to the PL ensemble size. (**C**) Correlation of chocolate seeking with the NAcore ensemble size. (**D**) Correlation of chocolate seeking to the PL ensemble size. (**E**) Correlation of cocaine seeking in dual conditioning with the NAcore ensemble size. (**F**) Correlation of cocaine seeking in dual conditioning to the PL ensemble size. Nucleus accumbens core (NAcore), prelimbic cortex (PL).

